# In Vitro Safety Signals for Potential Clinical Development of the Anti-Inflammatory Pregnane X Receptor Agonist FKK6

**DOI:** 10.1101/2023.10.21.563410

**Authors:** Zdeněk Dvořák, Barbora Vyhlídalová, Petra Pečinková, Hao Li, Pavel Anzenbacher, Alena Špičáková, Eva Anzenbacherová, Vimanda Chow, Jiabao Liu, Henry Krause, Derek Wilson, Tibor Berés, Petr Tarkowski, Dajun Chen, Sridhar Mani

## Abstract

**Background and purpose:** Based on the mimicry of microbial metabolites, functionalized indoles were demonstrated as the ligands and agonists of the pregnane X receptor (PXR). The lead indole, FKK6, displayed PXR-dependent protective effects in DSS-induced colitis in mice and *in vitro* cytokine-treated intestinal organoid cultures. Here, we performed the initial *in vitro* pharmacological profiling of FKK6.

**Experimental approach:** A complex series of cell-free and cell-based assays were employed. The organic synthesis, and advanced analytical chemistry methods were used.

**Key results:** FKK6-PXR interactions were characterized by hydrogen-deuterium exchange mass spectrometry. Screening FKK6 against potential cellular off-targets revealed high PXR selectivity. FKK6 has poor aqueous solubility but was highly soluble in simulated gastric and intestinal fluids. FKK6 was bound to plasma proteins and chemically stable in plasma. The partition coefficient of FKK6 was 2.70, and FKK6 moderately partitioned into red blood cells. In Caco2 cells, FKK6 displayed high permeability (A-B: 22.8 × 10^-6^ cm.s^-1^) and no active efflux. These data are indicative of essentially complete *in vivo* absorption of FKK6. FKK6 was rapidly metabolized by cytochromes P450, notably by CYP3A4 in human liver microsomes. Two oxidized FKK6 derivatives, including N6-oxide and C19-phenol, were detected, and these metabolites had 5-7 × lower potency as PXR agonists than FKK6. This implies that despite high intestinal absorption, FKK6 is rapidly eliminated by the liver, and its PXR effects are predicted to be predominantly in the intestines.

**Conclusion and implications:** The PXR ligand and agonist FKK6 has a suitable pharmacological profile supporting its potential preclinical development.

**BULLET POINT SUMMARY:** What is already known:

- Microbial metabolite mimic FKK6 is a hPXR agonist with anti-inflammatory properties in mice and human.
- The *in vitro* PXR binding, absorption, and metabolism have not been completely characterized.

What this study adds:

- PXR selectivity with unique binding mode, high intestinal cell permeability, rapid and complex microsomal metabolism.
- Initial testing for predicted metabolites shows reduced potency as PXR agonists.

Clinical significance:

- PXR effects of FKK6 are predicted to be predominantly in the intestines.
- FKK6 has a suitable pharmacological profile supporting its potential preclinical development.

## INTRODUCTION

Pregnane X receptor (PXR) is a nuclear receptor (*NR1I2*) that was initially discovered as a xenobiotic sensor and a culprit of drug-drug interactions through the induction of drug-metabolizing enzymes. Therefore, drug discovery suspended the substances that activated PXR from further testing. However, more recently, multiple physiological roles of PXR have been unveiled, including regulating inappropriate inflammation in various organs like the gastrointestinal tract (e.g., inflammatory bowel disease; IBD)[1–6]. Since the initial discovery of the PXR and its functional importance as a master regulator of the drug metabolism [7], a new pharmacotherapeutic paradigm might be important in that, for specific diseases (e.g., IBD), dialing-in PXR activity, especially in the intestines and not systemically, could safely expand the new drug space [8, 9]. Indeed, an example is the antibiotic rifaximin, which is used clinically in part for its PXR-agonist activity in treating inflammatory bowel disease [10]. Unlike rifaximin, which is non-absorbable from the intestine or formulations that are restricted to colon delivery only [11], many PXR-active drugs reach systemic circulation and cause adverse effects such as the induction of drug-metabolizing enzymes or perturbation of lipid and cholesterol metabolism. Existing PXR ligands also have substantial off-target toxicity [12]. In our prior work, we have established microbial metabolite mimicry as a novel strategy for drug discovery that allows exploiting previously unexplored parts of the chemical space [12]. We have reported functionalized indole derivatives as first-in-class non-cytotoxic PXR agonists as a proof of concept for microbial metabolite mimicry. The lead compound, FKK6 (Felix Kopp Kortagere 6), has been demonstrated as a PXR ligand and agonist that induced PXR-specific target genes *in vitro* and *in vivo* in mice. FKK6 displayed PXR-dependent protective effects in DSS-induced colitis in mice and human intestinal organoids stimulated with pro-inflammatory cytokines [12, 13]. In addition, FKK6 did not activate the aryl hydrocarbon receptor (AhR), which is a good sign of selectivity, given frequent dual agonist effects of indole derivatives, such as methylated indoles [14–16], microbial indoles [17, 18], substituted tryptamine [19] and imidazolopyridyl-modified FKK series [20]. Here, we carried out detailed *in vitro* pharmacological profiling of FKK6, using complementary sets of non-cellular and cell-based models, to assess FKK6’s suitability for potential preclinical development as a therapeutic for IBD. FKK6-PXR interactions were characterized by hydrogen-deuterium exchange mass spectrometry. Screening against several potential cellular off-targets revealed the high PXR-selectivity of FKK6. High solubility in intestinal simulated fluid, high *in vitro* permeability in Caco2 cells, no active efflux, and partition coefficient > 1, indicate essentially complete *in vivo* absorption of FKK6. The rapid clearance of FKK6 notably by CYP3A4, the formation of oxidized FKK6 metabolites with diminished PXR-agonist activity, and inhibitory effects of FKK6 against CYP3A4 imply that despite high intestinal absorption, it is unlikely to observe systemic PXR activation effects. We conclude that FKK6 has a suitable pharmacological profile supporting its potential preclinical development.

## MATERIALS AND METHODS

### Bottom-up Hydrogen Deuterium Exchange Mass Spectrometry (HDX-MS)

#### hPXR recombinant protein production

A pET28-MHL hPXR LBD-SRC1 expression plasmid was made by amplifying the hPXR LBD-SRC1 cassette from the pLIC vector with 15-bp extensions complementary to each end of the BsERI digested pET28-MHL backbone. The amplified DNA was inserted into the pET28-MHL vector by In-Fusion HD Cloning. Recombinant proteins were expressed as previously described [21]. Proteins were purified using of HisPur Ni-NTA resin (Thermo Fisher Scientific, 88222), followed by size exclusion FPLC (HiLoad 16/60 Superdex 200, GE Healthcare, Life Sciences) equilibrated with 150 mM NaCl, 50 mM HEPES (pH 8.2) and 0.5 mM DTT, at a flow rate of 1 mL/min.

To probe the protein-ligand interaction, bottom-up hydrogen-deuterium exchange mass spectrometry (HDX-MS) was employed. PXR protein was prepared at 10 μM in 20 mM HEPES, 150 mM NaCl pH 8.2, and 10% DMSO. A 1:10 protein:ligand ratio was used to probe the interactions, where the final concentration of the ligand was 100 μM with 1% DMSO. Samples were diluted into equilibration buffer (10 mM phosphate buffer, pH 7.5) or deuterated reaction buffer (10 mM phosphate buffer, 150 mM NaCl, pH 7.5 in D_2_O). The HDX reaction over labeling times of 1, 10, and 30 minutes at 25°C, and quenched with equal volumes of quench buffer (100 mM phosphate buffer, pH 2.5) at 0°C. An enzyme BEH Pepsin column (Cat#186007233) performed sample digestion at 15°C followed by peptide trapping and desalting on an ACQUITY UPLC BEH C18 VanGuard Pre-column (Cat#186003975). Digested peptides were separated using an ACQUITY UPLC BEH C18 column (Cat#186002346). All HDX experiments were conducted using M-class ACQUITY UPLC attached to Waters Cyclic IMS Mass Spectrometer. Peptide identification and uptake were analyzed using ProteinLynx Global server (PLGS) and DynanX. For protein illustration, PyMol 2.5 software was used.

### The activity of G protein-coupled receptors by PathHunter^®^ β-Arrestin assay

The PathHunter^®^ β-Arrestin assay monitors the activation of a GPCR in a homogenous, non-imaging assay format using a technology developed by DiscoveRx called Enzyme Fragment Complementation with β-galactosidase as the functional reporter. The enzyme is split into two inactive complementary portions (Enzyme Acceptor and Enzyme Donor) expressed as fusion proteins in the cell. Enzyme Acceptor is fused to β-Arrestin, and Enzyme Donor is fused to the GPCR of interest. When the GPCR is activated and β-Arrestin is recruited to the receptor, Enzyme Donor and Enzyme Acceptor complementation occurs, restoring β-galactosidase activity, which is measured using chemiluminescent PathHunter^®^ Detection Reagents. Calculations per Eurofins DiscoveRx^®^ Profiling Service recommendation suggest that the % activity for gpcrMAX agonist (> 30%) / antagonist (> 50%), and orphanMAX agonist (> 50%) is not indicative of interaction given the similarity in the limits posed by the mean and baseline RLU.

### Nuclear hormone receptor nhrMAX assay

PathHunter^®^ NHR Protein Interaction (Pro) and Nuclear Translocation (NT) assays monitor the activation of a nuclear hormone receptor in a homogenous, non-imaging assay format using a technology developed by DiscoveRx called Enzyme Fragment Complementation (EFC). The NHR Pro assay is based on the detection of protein-protein interactions between an activated, full-length NHR protein and a nuclear fusion protein containing Steroid Receptor Co-activator Peptide (SRCP) domains with one or more canonical LXXLL interaction motifs. The NHR is tagged with the ProLinkTM component of the EFC system, and the SRCP domain is fused to the Enzyme Acceptor component expressed in the nucleus. When bound by a ligand, the NHR will migrate to the nucleus and recruit the SRCP domain, whereby complementation occurs, generating a unit of active β-Galactosidase and production of a chemiluminescent signal. The NHR NT assay monitors the movement of a NHR between the cytoplasmic and nuclear compartments. The receptor is tagged with the ProLabel^TM^ component of the EFC system, and the Enzyme Acceptor is fused to a nuclear location sequence that restricts the expression of Enzyme Acceptor to the nucleus. Migration of the NHR to the nucleus results in complementation with Enzyme Acceptor generating a unit of active β-Galactosidase and production of a chemiluminescent signal. Calculations per Eurofins DiscoveRx^®^ Profiling Service recommendation suggest that the % activity for nhrMAX agonist (> 30%) / antagonist (> 50%) is indicative of potential interaction.

### The Comprehensive in Vitro Proarrhytmia Assay (CiPA) - IonChannelProfiler™

#### Human Sodium Ion Channel (hNa_V_1.5) Cell-Based QPatch Assay

Onset and steady-state block of peak hNa_V_1.5 current was measured using a pulse pattern, repeated every 5 sec, consisting of a hyperpolarizing pulse to -120 mV for a 200 ms duration, depolarization to -15 mV amplitude for a 40 ms duration, followed by step to 40 mV for 200 ms and finally a 100 ms ramp (1.2 V/s) to a holding potential of -80 mV. Peak current was measured during the step to -15 mV. Leak current was measured at the step pulse from -120 mV to -130 mV.

#### Human Calcium Ion Channel (hCa_V_1.2 L-type) Cell-Based QPatch Assay

After whole-cell configuration was achieved, a 50 ms step pulse from -60 mV to 10 mV in 15 s intervals for a total of three pulses was applied, and inward peak currents were measured upon depolarization of the cell membrane. Following the addition of the compound, five pulses of -80 mV to -60 mV for 5000 ms were applied before the three-step pulses.

#### Human Potassium Ion Channel (hK_V_11.1. hERG) Cell-Based QPatch Assay

After the whole cell configuration, the cell was held at -80 mV. A 500 ms pulse to -40 mV was delivered to measure the leaking current, subtracted from the tail current online. Then, the cell was depolarized to +40 mV for 500 ms and then to -80 mV over a 100 ms ramp to elicit the hERG tail current. This paradigm is delivered once every 8 s to monitor the current amplitude.

The peak current amplitude was calculated before and after compound addition for each ion channel. The amount of block was assessed by dividing the Test compound current amplitude by the Control current amplitude. Control is the mean respective ion channel current amplitude collected 15 s at the end of the control; Test Compound is the mean respective ion channel current amplitude collected in the presence of test compound at each concentration.

### Inhibition of cytochromes P450

Effects of FKK6 (10 μM) on the catalytic activities of drug-metabolizing cytochromes P450 were studied in human liver microsomes, and the quantitation of metabolites was carried out by HPLC-MS/MS techniques, as described elsewhere [22]. The experiments were performed at two independent research laboratories, i.e., Charles River Laboratories, Inc. (Wilmington, MA, USA) and Eurofins Discovery (Eurofins Panlabs, St. Charles, MO, USA). Substrates, their metabolites and specific control inhibitors, respectively, for each isoform were: CYP1A (phenacetin/acetaminophen; furafylline and fluvoxamine), CYP2B6 (bupropion / hydroxybupropion; clopridogel and ticlopidine), CYP2C8 (amodiaquine / desethylamodiaquine; montelukast and quercetin), CYP2C9 (diclofenac / 4’-hydroxydiclofenac; sulfaphenazole), CYP2C19 (omeprazole and S-mephenytoin / 5-hydroxyomeprazole and 4’-hydroxy-S-mephenytoin; oxybutynin and omeprazole), CYP2D6 (dextromethorphan/dextrorphan; quinidine), CYP3A (midazolam / 1-hydroxymidazolam; ketoconazole), CYP3A (testosterone / 6β-hydroxytestosterone; ketoconazole). Peak areas corresponding to the metabolite of each substrate were recorded. The percent of control activity was calculated by comparing the peak area obtained in the presence of the test compound to that obtained in the absence. The percent inhibition was calculated by subtracting the percent control activity from 100 for each compound.

### Inhibition of drug transporters

Effects of FKK6 (10 μM) on the activity of major human drug transporters was assessed by Eurofins facility, using techniques described elsewhere; briefly:

#### OCT1 and OCT2 (Organic Cation Transporter)

the activities were assessed by fluorimetry as an uptake of 4–4-dimethylaminostyryl-N-methylpyridinium in OCT1-CHO-K1 [23] and OCT2-CHO-K1 [24] recombinant cells. The reference inhibitor was verapamil (OCT1-IC_50_ = 7.5 μM; OCT2-IC_50_ = 35 μM).

#### OAT1 and OAT3 (Organic Anion Transporter)

the activities were assessed by fluorimetry as an uptake of 6-carboxyfluorescein in OAT1-CHO-K1 and OAT3-CHO-K1 recombinant cells [25]. The reference inhibitor was probenicid (OAT1-IC_50_ = 31 μM; OAT2-IC_50_ = 16 μM).

#### OATP1B1 and OATP1B3 (Organic Anion Transporting Polypeptide)

the activities were assessed by fluorimetry as an uptake of fluorescein-methotrexate in OATP1B1-CHO-K1 and OATP1B3-CHO-K1 recombinant cells [26]. The reference inhibitor was rifampicin (OATP1B1-IC_50_ = 3.4 μM; OATP1B3-IC_50_ = 3.9 μM).

#### MATE1 and MATE2-K (Multidrug And Toxin Extrusion protein)

the activities were assessed by fluorimetry as an uptake of 4′,6-diamidino-2-phenylindole in MATE1-HEK and MATE2-K-HEK recombinant cells [27]. The reference inhibitor was verapamil (MATE1-IC_50_ = 16 μM; MATE2-K-IC_50_ = 15 μM).

#### BCRP (Breast Cancer Resistance Protein)

the activity was assessed by fluorimetry as an uptake of Hoechst33342 in BCRP-CHO-K1 recombinant cells [28]. The reference inhibitor was KO143 (IC_50_ = 0.13 μM).

#### MRP2 (Multidrug Resistance-associated Protein)

the activity was assessed by fluorimetry as an uptake of 5(6)-carboxy-2’,7’-dichlorofluorescein in MRP2-HEK recombinant cells [29]. The reference inhibitor was MK571 (IC_50_ = 22 μM).

#### P-gp (Multidrug Resistance Protein 1)

the activity was assessed by fluorimetry as an uptake of calcein AM in MDR1-MDCKII recombinant cells [30]. The reference inhibitor was verapamil (IC_50_ = 5.6 μM).

#### BSEP (Bile Salt Export Pump)

the activity was assessed by scintillation counting as an uptake of ^3^H-taurocholic acid in BSEP-HEK membrane vesicles [31]. The reference inhibitor was cyclosporine A (IC_50_ = 0.54 μM).

### Aqueous solubility

Solubility was studied in PBS (pH 7.4) and simulated gastric and intestinal fluids using the shake-flask technique as described elsewhere [32]. The incubations were 2 h and 24 h at room temperature, and HPLC/UV-VIS determined the concentration of FKK6.

### Plasma proteins binding

A binding of FKK6 to human and mouse plasma proteins was studied by equilibrium dialysis (FKK6 – 10 μM; 4 h; 37°C) [33] and ultracentrifugation technique (FKK6 – 2 μM; 2.5 h; 37°C). The concentration of FKK6 was determined by HPLC/MS-MS. Acebutolol, quinidine, and warfarin were used as reference compounds. The peak areas of the FKK6 in the buffer (supernatant) (S_B_) and protein matrix (S_P_) were used to calculate percent binding according to the formula: protein binding % = 100×(S_P_ - S_B_)/ S_P_

### Plasma stability

FKK6 (2 μM) was incubated at 37°C with human or mouse plasma for 0 min, 10 min, 20 min, 30 min, 60 min, and 120 min. Propantheline (2 μM) was used as a reference compound that undergoes degradation in plasma.

### Partition coefficient

The standard OECD method No. 117 was used, based on the linear regression of partition coefficients of standards with the known log P values against the log of capacity factors of these compounds obtained by the HPLC analysis. Two different columns were used: (i) Agilent Poroshell 120 EC-18, 2.7 µm, 50 mm × 4.6 mm i.d.; (ii) ReproSil-Pur Basic C18, 5 µm, 200 mm × 4 mm i.d.

### Red blood cell partitioning

The assay was performed according to the standard protocols. The incubations with FKK6 (2 μM) or methazolamide (Met; 2 μM) were performed for 1 h at 37°C. The calculations were performed as follows (PL = plasma; H = hematocrit; WB = whole blood; C = concentration): *Adjusted Ratio*, which takes hematocrit into account: (C_spiked PL_ / C_PL from spiked WB_) × 1/(1-H) *Partition coefficient*: K_p(RBC/PL)_ = [(C_spiked PL_ / C_PL from spiked WB_) - 1] × (1/H) + 1 *% Partitioned* is the percentage of total compound not present in plasma from spiked WB. Red blood cell partitioning is indicated if the Adjusted Ratio is > 1.0, and if K_p(RBC/PL)_ > 0.

### Intrinsic clearance and CYP phenotyping

FKK6 (0.1 μM) was incubated at 37°C with human liver microsomes (HLM; 0.1 mg/mL) or human recombinant CYPs (CYP1A2, CYP2B6, CYP2C8, CYP2C9, CYP2C19, CYP2D6, CYP3A4) for the periods of 0 min, 15 min, 30 min, 45 min and 60 min, according to published protocols [34, 35]. HPLC/MS-MS determined the concentration of remaining parental FKK6 at individual time points. The half-life (t_1/2_) was estimated from the slope of the initial linear range of the logarithmic curve of the FKK6 remaining *vs* time, assuming the first-order kinetics. The apparent intrinsic clearance (CL_int_) was calculated according to the following formula: CL_int_ = 0.693/t_1/2_

### PXR reporter gene assay

Intestinal LS180 cells transiently transfected by lipofection (FuGENE^®^ HD Transfection Reagent) with pSG5-hPXR expression plasmid along with a luciferase reporter plasmid p3A4-luc were used for assessment of PXR transcriptional activity [12]. Cells were incubated for 24 h with tested compounds. Thereafter, the cells were lysed and luciferase activity was measured on the Tecan Infinite M200 Pro plate reader (Schoeller Instruments, Czech Republic). Incubations were carried out in quadruplicates (technical replicates), and the experiments were performed in three consecutive cell passages.

### Permeability assay

The assays were carried out in Caco-2 cells at 37°C [36]: A-B (0 min, 60 min); B-A (0 min, 40 min). HPLC-MS-MS determined concentrations of test compounds. The ***apparent permeability coefficient*** was calculated as follows: P_app_ (cm.s^-1^) = V_R_×C_R,end_ / Δt×A×(C_D,mid_ – C_R,mid_), where V_R_ is the volume of the receiver chamber; C_R,end_ is the concentration of the test compound in the receiver chamber at the end time point, Δt is the incubation time and A is the surface area of the cell monolayer;. C_D,mid_ is the calculated mid-point concentration of the test compound in the donor side, which is the mean value of the donor concentration at time 0 minute and the donor concentration at the end time point; C_R,mid_ is the mid-point concentration of the test compound in the receiver side, which is one half of the receiver concentration at the end time point. The ***recovery*** of the test compound was calculated as follows:

Recovery % = 100×(V_D_×C_D,end_ + V_R_×C_R,end_)/V_D_×C_D0_, where V_D_ and V_R_ are the volumes of the donor and receiver chambers, respectively; C_D,end_ (C_R,end_) is the concentration of the test compound in the donor (receiver) sample at the end time point; C_D0_ is the concentration of the test compound in the donor sample at time zero. Fluorescein was used as the cell monolayer integrity marker. Fluorescein permeability assessment (in the A-B direction at pH 7.4 on both sides) was performed after the permeability assay for the test compound. The cell monolayer that had a fluorescein permeability of less than 1.5 × 10^-6^ cm.s^-1^ was considered intact.

### *In silico* prediction and synthesis of FKK6 metabolites

The putative oxidized metabolites of FKK6, including DC73 (N-oxide; atom N6) and DC97 (phenol; atom C19), were predicted with high probability by using FAst MEtabolizer software FAME2 (Figure S2) [37]. Both compounds were synthesized as follows:

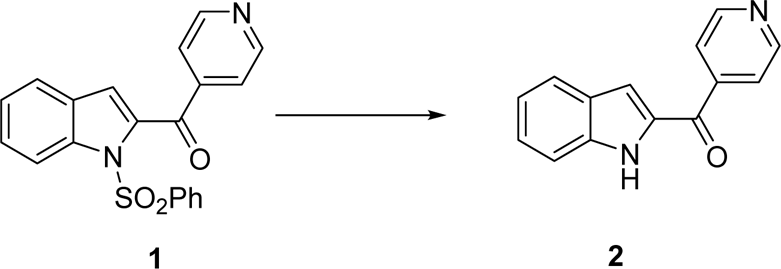

Synthesis of (1H-indol-2-yl)(pyridin-4-yl)methanone **(2)**: (1-(phenylsulfonyl)-1H-indol-2-yl)(pyridine-4-yl)methanone **(1)** (278 mg, 1 eq, 767 µmol, prepared by the reference method [12]), N,N,N-trimethylhexadecan-1-aminium bromide (14.0 mg, 0.05 eq, 38.4 µmol), potassium hydroxide (215 mg, 5 eq, 3.84 µmol) were charged to a microwave vial. THF (1.5 mL) and water (0.5 mL) were then added. The reaction mixture was heated at 70 °C by microwave for 3 hrs. After cooling to room temperature, water (5 mL) was added to the mixture, followed by EtOAc (3 mL × 3) for extraction. The combined organic phases were dried over Na_2_SO_4_, concentrated by vacuum, and purified by chromatography on silica gel (0 -10% MeOH in DCM) to afford compound **(2)** for 119 mg (yield: 70%). NMR spectra were consistent with reference data.

Synthesis of (1-((phenyl-d5)sulfonyl)-1H-indol-2-yl)(pyridin-4-yl)methanone **(3)** [38]: To a stirred mixture of (1-(phenylsulfonyl)-1H-indol-2-yl)(pyridine-4-yl)methanone (48 mg, 1 eq, 0.22 mmol), tetrabutylammonium bromide (21 mg, 0.3 eq, µmol) in benzene (2 mL), was added 1N aq. NaOH (0.66 mL, 3 eq, 0.66 mmol). The reaction mixture turned brown-red. Benzylsulfonyl chloride d-5 (59 mg, 1.5 eq, 0.32 mmol, prepared by reference method) in benzene (0.3 mL) was then added. The reaction mixture was stirred quickly for 5 minutes to achieve complete conversion as indicated by TLC. Water (2 mL) was added to the mixture, followed by EtOAc (5 mL × 2) for extraction. The combined organic phases were dried over Na_2_SO_4_, concentrated by vacuum, and purified by chromatography on silica gel (0 -10% MeOH in DCM) to afford compound **(3)** for 43 mg (yield: 65 %). HRMS: calculated 367.1039, found 368.1111 [M+1]^+^

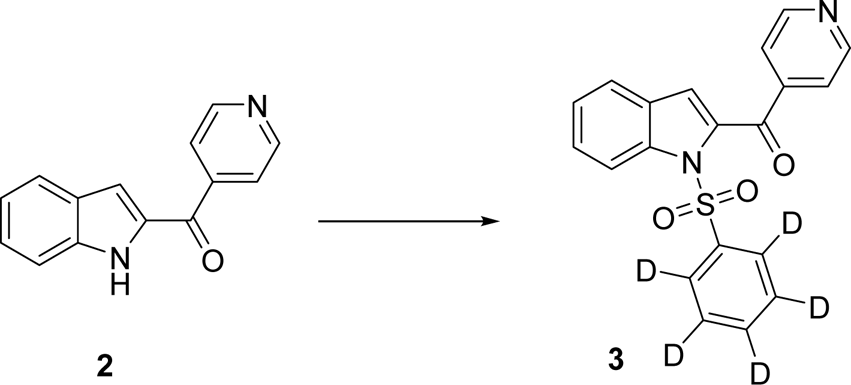

Synthesis of 1-(1-((phenyl-d5)sulfonyl)-1H-indol-2-yl)-1-(pyridin-4-yl)but-3-yn-1-ol **(4)** This synthesis was achieved by the same method like the non-deuterated version as described in the reference [12].

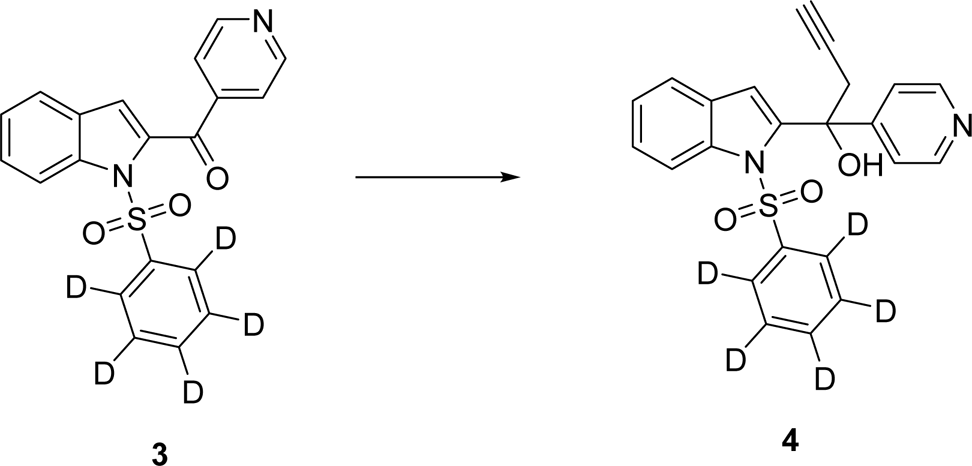

Synthesis of 4-(1-hydroxy-1-(1-(phenylsulfonyl)-1H-indol-2-yl)but-3-yn-1-yl)pyridine 1-oxide **(6) (DC73)**: To a solution of 1-(1-(phenylsulfonyl)-1H-indol-2-yl)-(1-pyridin-4-yl)-but-3-yn-1-ol (48.9 mg, 1 eq, 121 µmol, synthesized by the reference method [12]) in DCM (1.5 mL), was added meta-chloroperoxybenzoic acid (43.8 mg, 77% Wt, 1.6 eq, 195 µmol) in one portion. The reaction mixture was stirred overnight. The reaction mixture was filtered, the filtrate was washed with aq. Na_2_S_2_O_3_, dried over Na_2_SO_4_, concentrated in vacuo and purified by flash chromatography on silica gel (30-90% acetone in DCM) to afford the desired product as solid (27 mg, yield 53%).

^1^H-NMR (300 mHz, CDCl_3_) δ_H_ 8.11 (d, *J* = 8.4 Hz, 1H), 8.03 (d, *J* = 7.3 Hz, 2H), 7.73-7.62 (m, 4H), 7.51-7.46 (m, 2H), 7.43-7.37 (m, 1H), 7.35-7.29 (m, 4H), 5.61 (br, 1H), 3.37-3.23 (m, 2H), 2.51 (t, *J* = 2.6 Hz, 1H)

^13^C-NMR (150 mHz, CDCl_3_) δ_C_ 142.9, 141.2, 138.4, 138.3, 138.0, 134.2, 129.3, 127.9, 126.4, 125.8, 124.6, 123.7,121.8, 115.2, 114,4, 78.8, 74.3, 73.3, 35.0

ESI-MS: 419.1 [M+1]^+^

HRMS: calculated: 418.0987; observed: 419.1053 [M+1]^+^

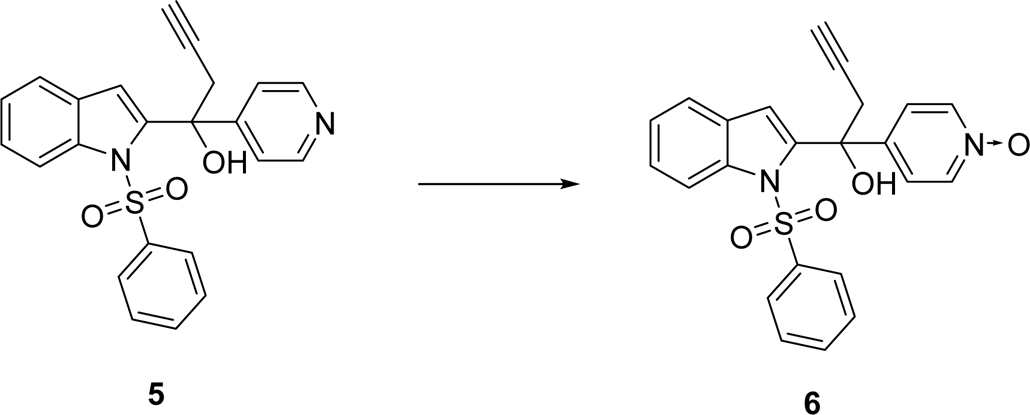

Synthesis of 4-((2-(1-hydroxy-1-(pyridin-4-yl)but-3-yn-1-yl)-1H-indol-1-yl)sulfonyl)phenol (9) (DC97):

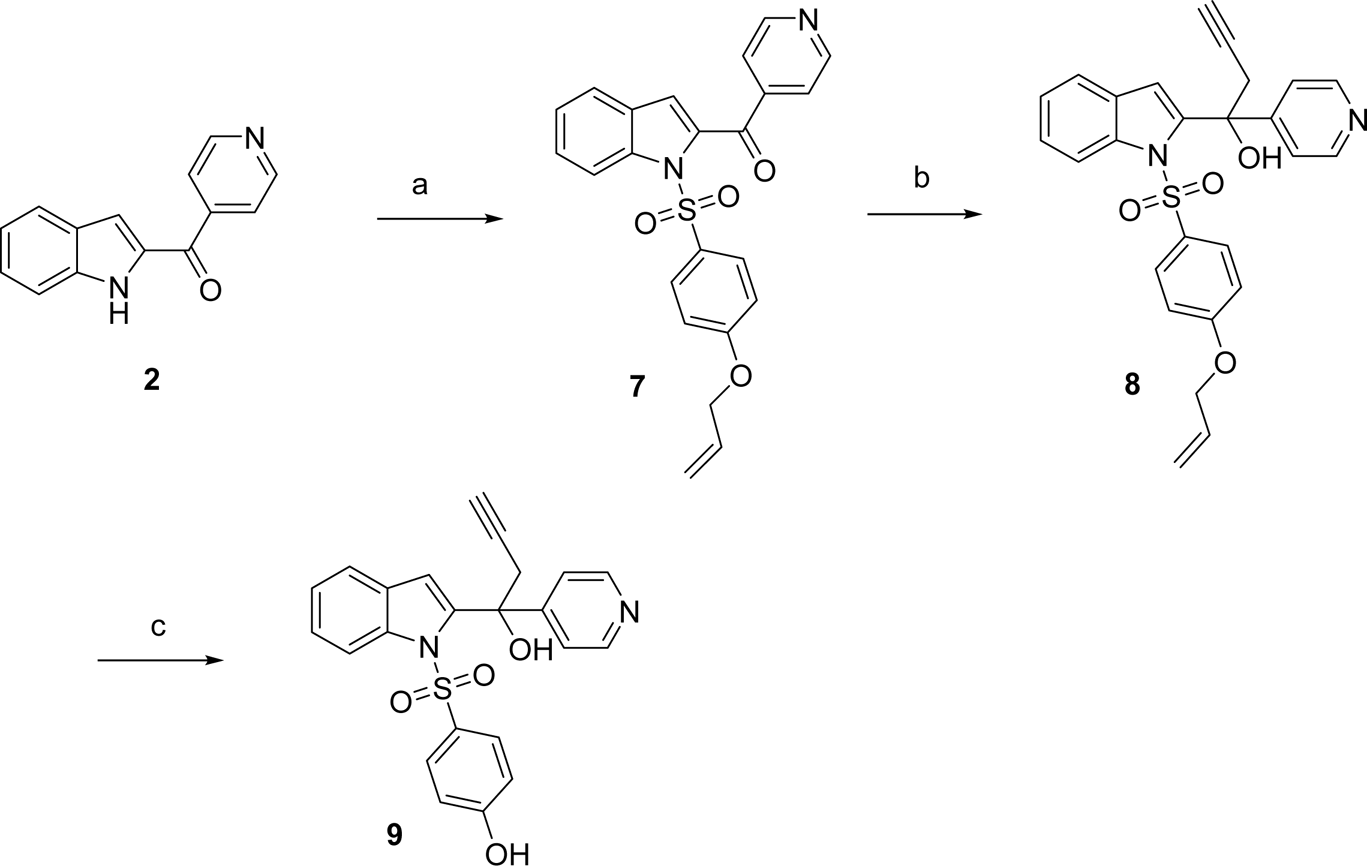

### Reaction Conditions

a. aq. NaOH, Bu_4_NBr(cat), Benzene, 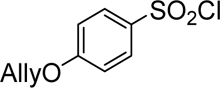 (purchased from Enamine.net)
b. 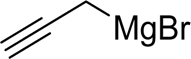 THF
c. K_2_CO_3_, Pd(PPh_3_)_4_, MeOH

Compound **(7)**, **(8)** were synthesized by the methods like compound **(3)**, **(4).** A solution of 1-(1-((4-(allyloxy)phenyl)sulfonyl)-1H-indol-2-yl)-1-(pyridin-4-yl)but-3-yn-1-ol (compound **(8)**, 41 mg, 1 Eq, 89 µmol), Pd(Ph_3_P)_4_ (2.1 mg, 0.02 Eq, 1.8 µmol) in MeOH (1 mL) was stirred at rt under N_2_ for 5 min. K_2_CO_3_ (50 mg, 4.0 Eq, 0.36 mmol) was then added. The flask was sealed to remove air by vacuum, and flushed with N_2_The reaction mixture was stirred for 4 hrs. Aqueous NH_4_Cl (1 mL) was added to quench the reaction, followed by extraction with DCM (3 mL × 3). The combined organic phases were dried over Na_2_SO_4_, concentrated, and purified by flash chromatography on silica gel (35-60% EtOAc in Hexane) to afford compound **(9)** as solid (17 mg, yield 35%).

^1^H-NMR (300 mHz, CDCl_3_) of compound **7**:

δ_H_ 8.83 (dd, *J* = 4.5, 1.5 Hz, 2H), 8.12 (d, *J* = 8.5 Hz, 1H), 7.93-7.88 (m, 2H), 7.75 (dd, *J* = 4.4, 1.6 Hz, 2H), 7.58 (d, *J* = 7.9 Hz, 1H), 7.51-7.46 (m, 1H), 7.33-7.28 (m, 1H), 7.02 (s, 1H), 6.96 (dt, *J* = 7.1, 1.9 Hz, 2H). 6.02-5.93 (m, 1H), 5.37 (dd, *J* = 17.3, 1.4 Hz, 1H), 5.40-5.27 (m, 2H), 4.53 (dt, *J* = 5.3, 1.3 Hz, 2H)

MS (ESI) of compound **7**: 419.1 [M+1]^+^

^1^H-NMR (300 mHz, CDCl_3_) of compound **8**: δ_H_ 8.49 (d, *J* = 5.8 Hz, 2H), 8.03 (d, *J* = 8.2 Hz, 1H), 7.60-7.57 (m, 1H), 7.37-7.24 (m, 6H), 7.13 (s, 1H), 6.74-6.69 (m, 2H), 6.01-5.89 (m, 1H), 5.39-5.26 (m, 2H), 4.48 (dt, *J* = 5.3, 1.3 Hz, 2H), 3.12 (d, *J* = 2.5 Hz, 2H), 2.08 (t, *J* = 2.6 Hz, 1H)

MS (ESI) of compound **8**: 459.1 [M+1]^+^

^1^H-NMR (300 mHz, CDCl_3_) of compound **9:**

δ_H_ 8.27-8.25 (m, 3H), 7.66-7.30 (m, 5H), 7.18 (s, 1H), 6.89 (d, *J* = 8.9 Hz, 2H), 6.60 (d, *J* = 8.9 Hz, 2H), 5,75 (br, 1H), 3.13-2.99 (m, 2H), 2.05 (t, *J* = 2.6 Hz, 1H)

^13^C-NMR (150 mHz, CDCl_3_) δ_C_ 163.3, 154.5,147.3, 141.0, 138.4, 132.1,128.6, 128.0, 126.1, 124.2, 122.2, 121.6, 115.8, 115.4, 113.1, 78.9, 74.7, 72.9, 35.7

ESI-MS of compound **9:** 419.1 [M+1]^+^

HRMS of compound **9**: calculated: 418.0987; observed: 419.1053 [M+1]^+^

### FKK6 metabolism by human liver microsomes

Microsomal fraction of human liver homogenate (HLM, human liver microsomes) was obtained from Advancell (Barcelona, Spain). HLM were prepared by differential centrifugation using standard method and kept frozen until used. Incubation mixture contained 50 mM potassium phosphate buffer (pH 7.4), NADPH-generating system (0.5 mM NADP^+^, 3.8 mM isocitrate, 0.09 unit/mL of isocitrate dehydrogenase and 5 mM MgCl_2_), pooled HLM (130 pmol/reaction mixture) and the FKK6 (50 μM) in total volume of 250 μL. The reaction was stopped after 30 min incubation at 37 °C with two volumes of methanol. The reaction mixture was centrifuged at 18 400 g and the supernatant was analyzed for potential metabolites. HPLC system Shimadzu Prominence (Shimadzu, Kyoto, Japan) was used with Chromolith 100 (5 μm) RP-18 endcapped column (Merck, Darmstadt, Germany), mobile phase was 60% methanol: 40% deionized water, flow rate 0.6 ml.min^-1^), with UV detection at 254 nm. To prove an involvement of P450 in the formation of metabolites, two approaches were applied: (i) the reaction mixture was bubbled for 2 min thoroughly by carbon monoxide, which binds strongly to the P450 heme iron making the reaction impossible; (ii) an experiment with heat-inactivated microsomes (30 min) in a thermostated water bath (60 °C) was performed to inactivate the P450 enzymes.

### Identification of FKK6 microsomal metabolites by LC/MS

The identification and quantification of FKK6 metabolites was carried out *via* UHPLC-HESI-MS/MS-PDA technique, using a liquid chromatograph (Thermo UltiMate™ 3000, Thermo Fisher Scientific, Waltham, MA, USA) with triple quadrupole mass spectrometer (TSQ Quantum Access Max, Thermo Fisher Scientific, Waltham, MA, USA). Separation of compounds (FKK6, metabolites DC97 and DC73) was achieved *via* isocratic elution using an Acquity UPLC® BEH (100 × 2.1 mm; 1.7 µm; Waters, Milford, MA, USA) chromatographic column at a flow rate of 0.25 mL.min^-1^ using a mobile phase consisting of 65% of 15 mM ammonium formate, pH 3.0 (A) and 35% acetonitrile (B). The injected volume was 5 μL. The chromatographic column was kept at 40 °C. Collision spectra were recorded using a product scan mode (scan range *m/z* 50 – 450 parent mass 419, collision energy 50 eV), and the ionization was carried out at spray voltage 3 kV, vaporizer and capillary temperatures 300°C and 320°C respectively. Sheath, ion sweep, and auxiliary gas (nitrogen) pressures were set to 45, 1, and 15 psi, respectively. The MS/MS and PDA spectra were compared with the authentic standards.

## RESULTS

### Hydrogen-deuterium exchange mass spectrometry (HDX-MS) of FKK6-PXR

HDX-MS may provide insight into ligand binding interactions that are not accessible by classical structural methods, such as X-ray crystallography. HDX-MS confirmed the binding interactions between PXR and SR12813, a known PXR agonist. In the interest of discovering another agonist (FKK6), the rate of deuterium uptake was compared between ligand-free, SR12813-bound and FKK6-bound PXR. Figure 1 displays the regions identified by HDX, with a sequence coverage of 70.6% and redundancy for covered amino acids of 2.24 (Figure S1). In the presence of SR12813 and FKK6, peptides 293-307 and 319-335 showed similar conformational changes. The observed summed differences in deuterium for peptide 293-307 and 319-335 suggests a weak conformational change in the presence of FKK6 (-5% < x < - 10%), whereas a larger conformational change occurred in the presence of SR12813 (difference in deuterium uptake > -15%). Interestingly, extracted from the HDX-MS analysis, there are two different conformational changes; in the presence of SR12813, helix α3 (peptide 238-257) and α10 (peptide 397-410) stabilizes. Helix α5 (peptide 276-288) becomes destabilized, allowing for more deuterium exchange in the presence of FKK6.

**Figure 1.**
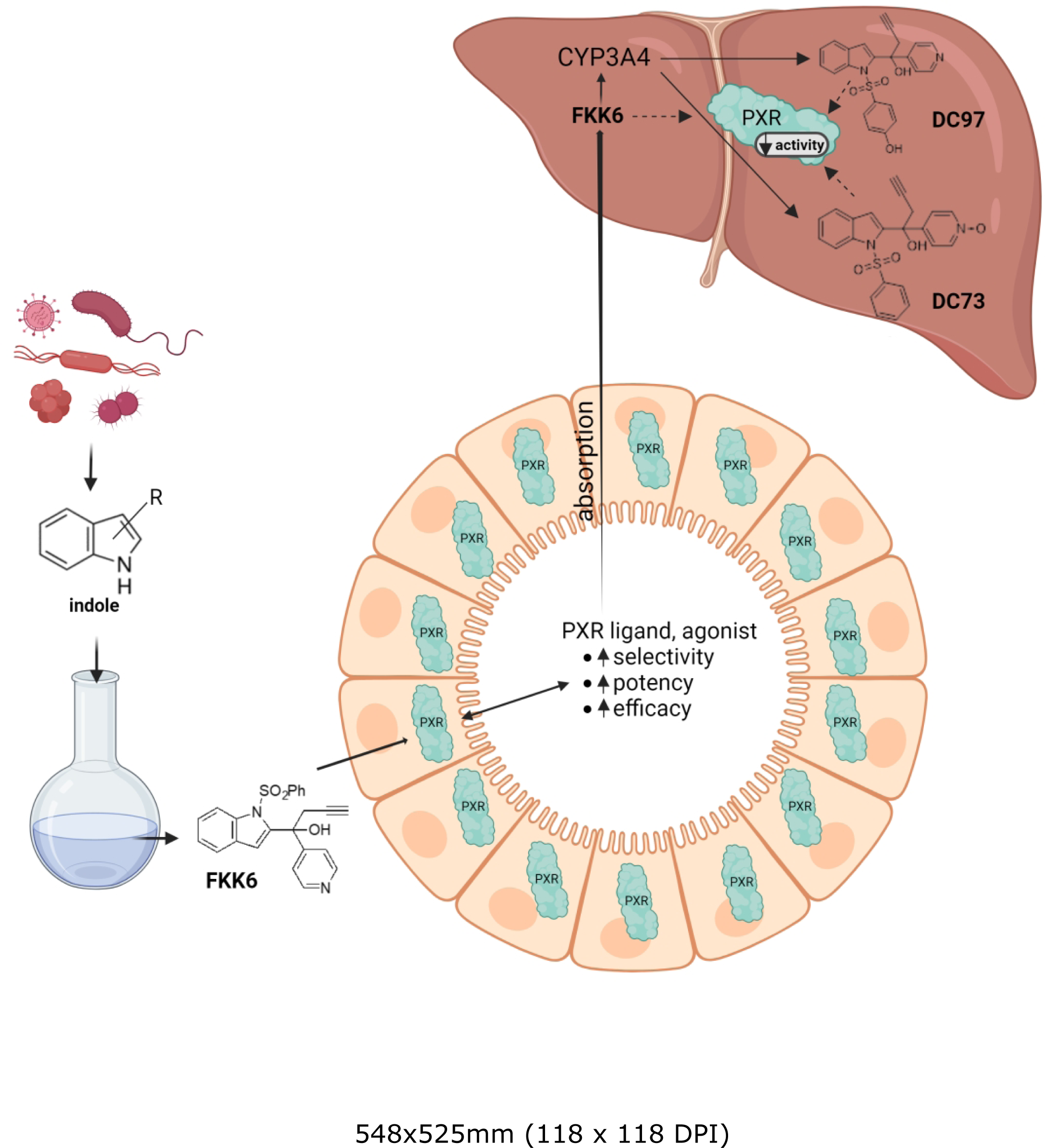
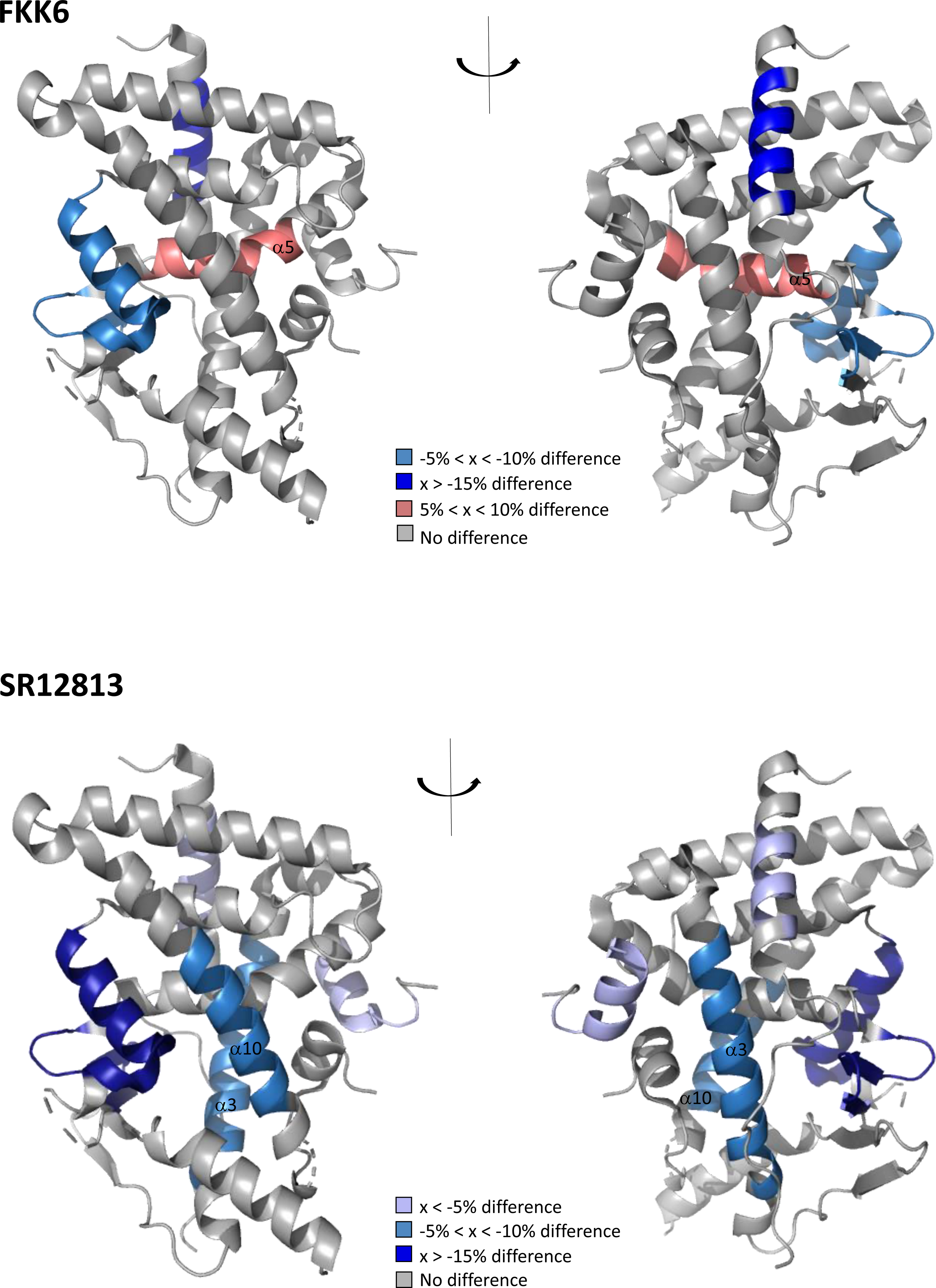
Visual representations of the differential hydrogen-deuterium exchange analysis of PXR binding interactions with SR12813 and FKK6 (PDB code 3HVL). The relative fractional difference in percentage of deuterium incorporation (%D) were summed and color coded accordingly on the visual structures. The dark blue denotes differences >-15%, light blue (-5%<x<-10%), where the complex (protein-ligand) exchanges less deuterium. The red denotes conformational changes where the complex incorporates more deuterium.

### FKK6 off-targets counter-screen

Whereas FKK6 is deemed a PXR-selective potential drug for treating intestinal inflammatory diseases, we performed a complex counter-screen to identify possible common pharmacological off-targets. First, we studied the effects of FKK6 (10 µM) on G protein-coupled receptors (GPCR), including gpcrMAX Panel (168 receptors; agonist and antagonist) and orphanMAX Panel (73 receptors; agonist). The activity of GPCRs was monitored by the PathHunter^®^ β-Arrestin assay, a technology developed by Eurofins DiscoveRx^®^. FKK6 did not activate any 241 GPCRs tested or antagonized any 168 non-orphan GPCRs (Figure 2A). Since PXR is a nuclear receptor, we also examined the effects of FKK6 on a panel of 19 steroid and nuclear receptors, using PathHunter^®^ NHR Protein Interaction and Nuclear Translocation assays. None of the examined receptors was activated or antagonized by FKK6 (Figure 2B). By using the Comprehensive in Vitro Proarrhytmia Assay CiPA - IonChannelProfiler™, we show that FKK6 does not block sodium (*hNa_V_1.5*), potassium (*hK_V_11.1. hERG*), and calcium (*hCa_V_1.2 L-type*) ion channels (Figure 2C). The effects of FKK6 on membrane drug transporters were investigated in a series of recombinant cell lines as previously described [23–31]. The activity of Bile Salt Export Pump (BSEP), Organic Anion Transporter 1B1 (OATP1B1), and Organic Anion Transporter 1B3 (OATP1B3) was inhibited by FKK6 (10 µM) down to 44%, 34% and 36% of control activity, respectively. The activity of Breast Cancer Resistance Protein (BCRP), Multidrug And Toxin Extrusion proteins (MATE1,2), Multidrug Resistance-associated Protein (MRP2), Organic Anion Transporters (OAT1,3), Organic Cation Transporters (OCT1,2), and Multidrug Resistance Protein 1 (P-gp) was not inhibited by FKK6 (Figure 2D). The capability of FKK6 to inhibit xenobiotic-metabolizing cytochromes P450 was studied in human liver microsomes. Surprisingly, FKK6 (10 µM) caused very strong inhibition of CYP2B6, CYP2C8, CYP2C9, CYP2C19, CYP2D6 and CYP3A4, whereas it did not inhibit CYP1A (Figure 2E). The data were acquired from two independent research laboratories. The inhibitory profile of FKK6, a non-specific pan-inhibitor, suggests it might directly interact with a common enzyme component, such as a heme in the active site.

**Figure 2.**
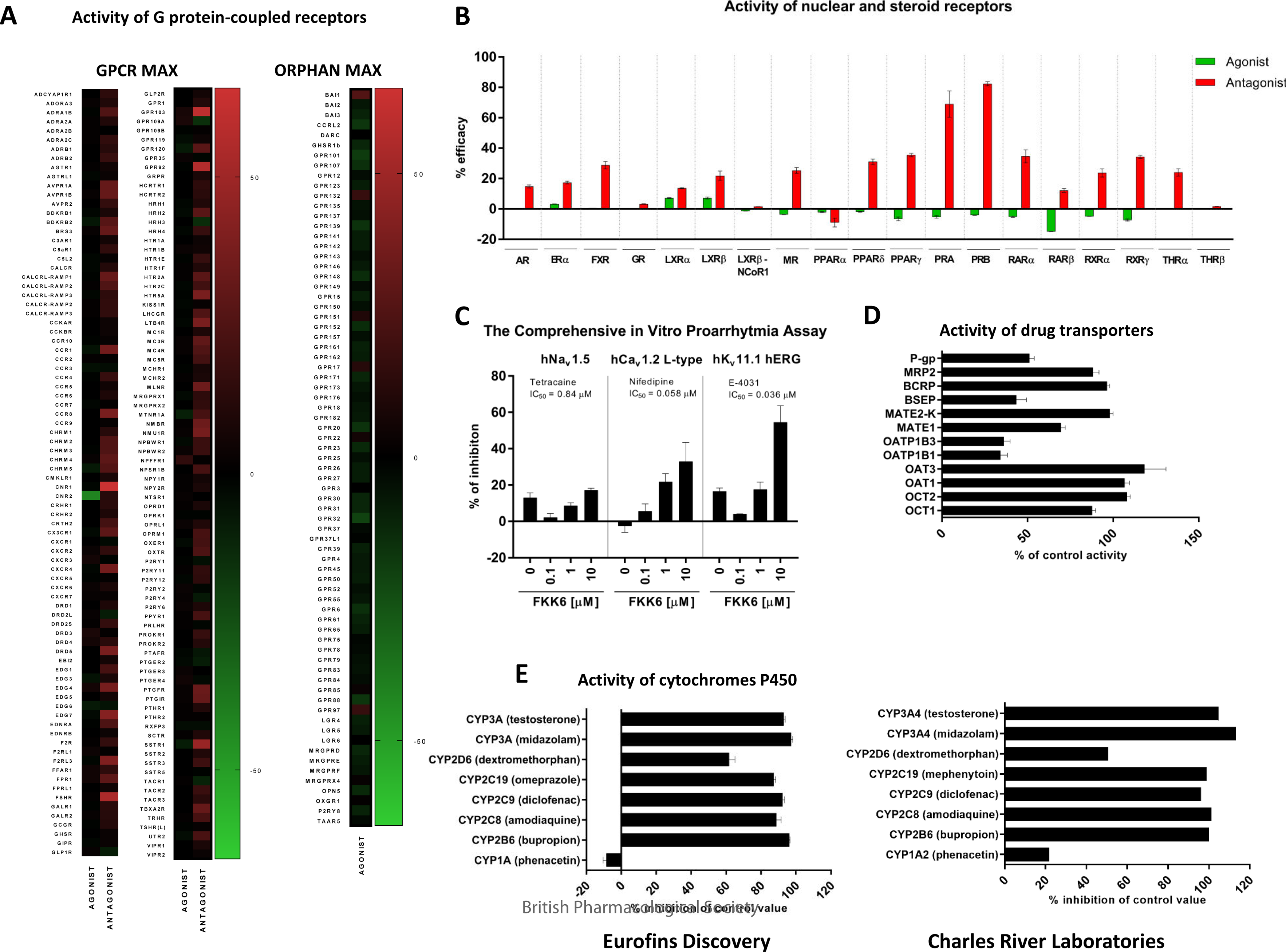
FKK6 off-targets counter-screen. ***(A) Activity of G protein-coupled receptors by PathHunter^®^ β-Arrestin assays.*** Assays with FKK6 (10 μM) (gpcrMAX Panel; orphanMAX Panel) were performed in duplicate, and mean values are presented. *<u>AGONIST mode:</u>* Double-gradient color map depicts percentage difference from control basal activity of the receptor; (+)-red values correspond to activation (agonism), (-)-green values correspond to inhibition of basal activity (inverse agonism); *<u>ANTAGONIST mode:</u>* Double-gradient color map depicts percentage difference from receptor activity achieved by model agonist in concentration of EC_80_; (+)-red values correspond to inhibition of agonist activity (antagonism), (-)-green values correspond to potentiation of agonist activity. ***(B) Activity of nuclear and steroid receptors.*** Bar graph shows agonist (green) and antagonist (red) effects of 10 μM FKK6 on the panel of nuclear and steroid receptors. Assays were performed in duplicate and mean values are presented. ***(C) The Comprehensive in Vitro Proarrhytmia Assay CiPA - IonChannelProfiler™***. Effects of FKK6 (0.1 μM; 1 μM; 10 μM) on the activity of sodium (*hNa_V_1.5*), potassium (*hK_V_11.1. hERG*) and calcium (*hCa_V_1.2 L-type*) ion channels. Data are expressed as percent inhibition relative to control. The IC_50_ values for reference antagonists are inserted in the graph. ***(D) Inhibition of drug transporters.*** The bar graph shows effects of 10 μM FKK6 on the activity of selected drug transporters. Data are expressed as a percentage of control activity in the absence of test compound. ***(E) Inhibition of cytochromes P450.*** Bar graph shows effects of 10 μM FKK6 on the activity of drug-metabolizing cytochromes P450. Data are expressed as percentage of inhibition as compared to control values.

### *In vitro* pharmacology of FKK6

In the next series of experiments, we characterized the pharmacological properties of FKK6 using a complementary set of *in vitro* cell-free and cell-based models.

The aqueous solubility of FKK6 was studied by using the shake-flask technique. Whereas FKK6 was nearly insoluble in PBS (pH 7.4), it displayed high solubility (150 µM – 200 µM) in simulated gastric (pH 1.2) and intestinal (pH 6.8) fluids (Figure 3A). These data might indicate strong pH-dependent bio-solubility of FKK6. Using equilibrium dialysis and ultracentrifugation techniques, we found that a high portion of FKK6 (96%-99 %) is bound to human and mouse plasma proteins (Figure 3B). The plasmatic stability of FKK6 (2 μM) was relatively high, yielding 79% and 87% of initial levels after 120 min of incubation with human and mouse plasma, respectively (Figure 3C).

**Figure 3.**
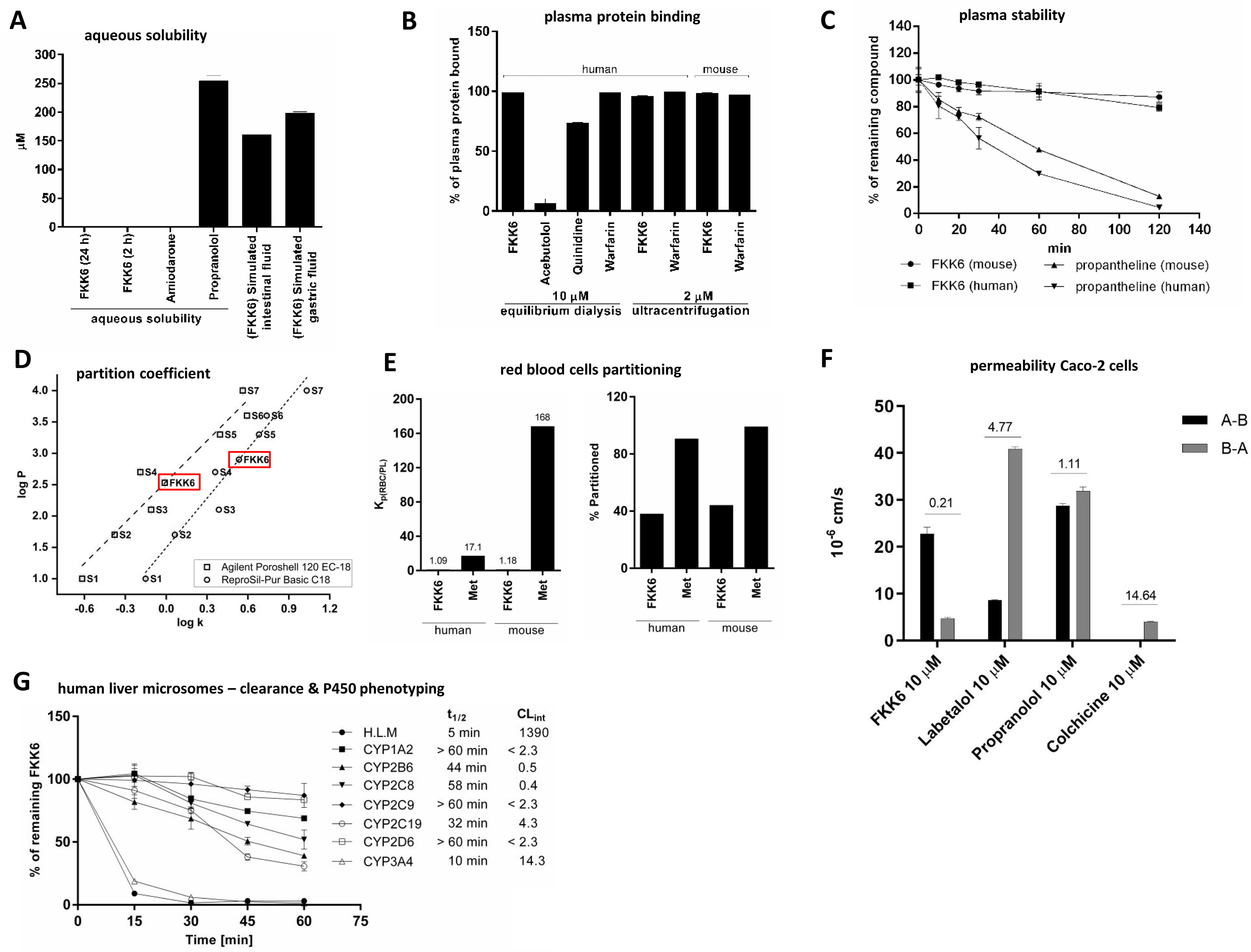
*In vitro* pharmacology of FKK6. ***(A) Aqueous solubility.*** The incubations were 2 h and 24 h at room temperature. ***(B) Binding to plasma proteins.*** A binding of FKK6 and reference compounds to human and mouse plasma proteins was studied by equilibrium dialysis and ultracentrifugation. ***(C) Plasma stability.*** A stability of FKK6 (2 μM) and reference compound propantheline was determined in human and mouse plasma in incubation time up to 120 min. ***(D) Partition coefficient.*** Log P (FKK6) was determined according to the standard OECD method No.117, using two different HPLC columns. ***(E) Red blood cells partitioning.*** Incubations with FKK6 (2 μM) or methazolamide (Met; 2 μM) were performed for 1 h at 37°C. Red blood cell partition coefficient (K_p(RBC/PL)_) was calculated as described in methods section. ***(F) Permeability in Caco-2 cells.*** The assays with FKK6 (10 μM) and reference compounds were carried out at 37°C, with end-point times 60 min (A-B) and 40 min (B-A). Inserted are values of efflux ratios. ***(G) Intrinsic clearance and CYP phenotyping.*** FKK6 (0.1 μM) was incubated at 37°C with human liver microsomes (HLM) or human recombinant CYPs for 0 min, 15 min, 30 min, 45 min and 60 min. The concentration of remaining parental FKK6 at individual time points was determined by HPLC/MS-MS, and the half-life (t_1/2_) and apparent intrinsic clearance (CL_int_) were calculated.

The partition coefficient (log P) of FKK6 was determined as 2.54 and 2.86 by standard OECD method No. 117 using two different HPLC columns (Figure 3D). Using red blood cell partitioning assay, we determined the red blood cell partition coefficient (K_p(RBC/PL)_) of FKK6 to be 1.09 and 1.18 in human and mouse blood, respectively. Accordingly, the percentage of FKK6 partitioned into human and mouse red blood cells was 38% and 44%, respectively (Figure 3E). These data reveal a moderate partition of FKK6 into red blood cells. The intestinal permeability and potential bioavailability of FKK6 were assessed in a standard model of differentiated polarized Caco2 cells. The absorptive (A-B) and secretory (B-A) apparent permeability coefficient P_app_ of FKK6 was 22.8 × 10^-6^ cm.s^-1^ and 4.7 × 10^-6^ cm.s^-1^, respectively, which indicates that FKK6 is highly permeable compound. Colchicine (P_app(A-B)_ 0.3 × 10^-6^ cm.s^-1^) and propranolol (P_app(A-B)_ 28.8 × 10^-6^ cm.s^-1^) were used as the reference low- and high-permeable compounds, respectively (Figure 3F). The efflux ratio (P_app(B-A)_ / P_app(A-B)_) of FKK6 was ∼ 0.2, which indicates that FKK6 is not a subject of active efflux. The actively effluxed compounds labetalol and colchicine displayed efflux ratios ∼ 14 and ∼ 5, respectively (Figure 3F). The recovery of FKK6 was about 60% (data not shown), which was comparable with test compounds. Therefore, the permeability data should not be underestimated. Overall, high *in vitro* permeability of FKK6 in Caco2 cells (with no active efflux), the partition coefficient of FKK6 > 1, and high FKK6 solubility in the intestinal simulated fluid is correlated with essentially complete *in vivo* absorbance of FKK6.

The intrinsic clearance of FKK6 was studied in human liver microsomes. The FKK6 was metabolized extensively with a half-life (t_1/2_) of 5 min, and the apparent intrinsic *in vitro* clearance (CL_int_) was 1390 μL/min/mg. The relative contribution of individual isoforms of cytochromes P450 on FKK6 hepatic clearance was investigated by using recombinant P450s (phenotyping). The most rapid disappearance of FKK6 was observed with CYP3A4, having t_1/2_ ∼ 10 min (Figure 3G). The high clearance of FKK6 may be a limiting factor for bioavailability, as it may undergo first-pass metabolism in the liver after oral dosing, which can limit the amount of drug reaching the circulation, even if the compound is well absorbed.

### *In vitro* metabolism of FKK6

The data from human liver microsomes (HLM) indicated that cytochromes P450, notably CYP3A4, rapidly metabolizes FKK6. Therefore, we used FAst MEtabolizer software FAME2 to predict *in silico* putative metabolites of FKK6. Two oxidized FKK6 derivatives, including DC73 (N-oxide; atom N6) and DC97 (phenol; atom C19), were predicted with high probability (Figure S2) and synthesized *de novo* (ref.: Methods section). Using reporter gene assay in transiently transfected intestinal LS180 cells, we observed that both FKK6 putative metabolites DC73 and DC97 dose-dependently activate PXR. The efficacies of DC73 and DC97 were like that of FKK6. In contrast, the potencies of DC73 (EC_50_ 7.8 µM) and DC97 (EC_50_ 5.1 µM) were 5-7-fold lower as compared to that of FKK6 (EC_50_ 1.2 µM) (Figure 4A). The metabolism of FKK6 in clearance and phenotyping experiments was assessed indirectly as a decrease of parental FKK6 in the reaction mixture with HLM. Therefore, we incubated FKK6 in high concentration (50 µM) with human liver microsomes, and we detected two new peaks by UV/HPLC, compared to control HLM mixtures. The retention time of parental FKK6 in the Reversed Phase HPLC system was 8.9 min. In contrast, putative FKK6 metabolites had retention times 5.1 min (M1) and 5.7 min (M2), suggesting they have higher polarity than parental FKK6. Saturation of HLM with carbon monoxide CO and inactivation of HLM by a heat abrogated FKK6 metabolism, verifying cytochrome P450-dependent formation of M1 and M2 metabolites (Figure 4B).

**Figure 4.**
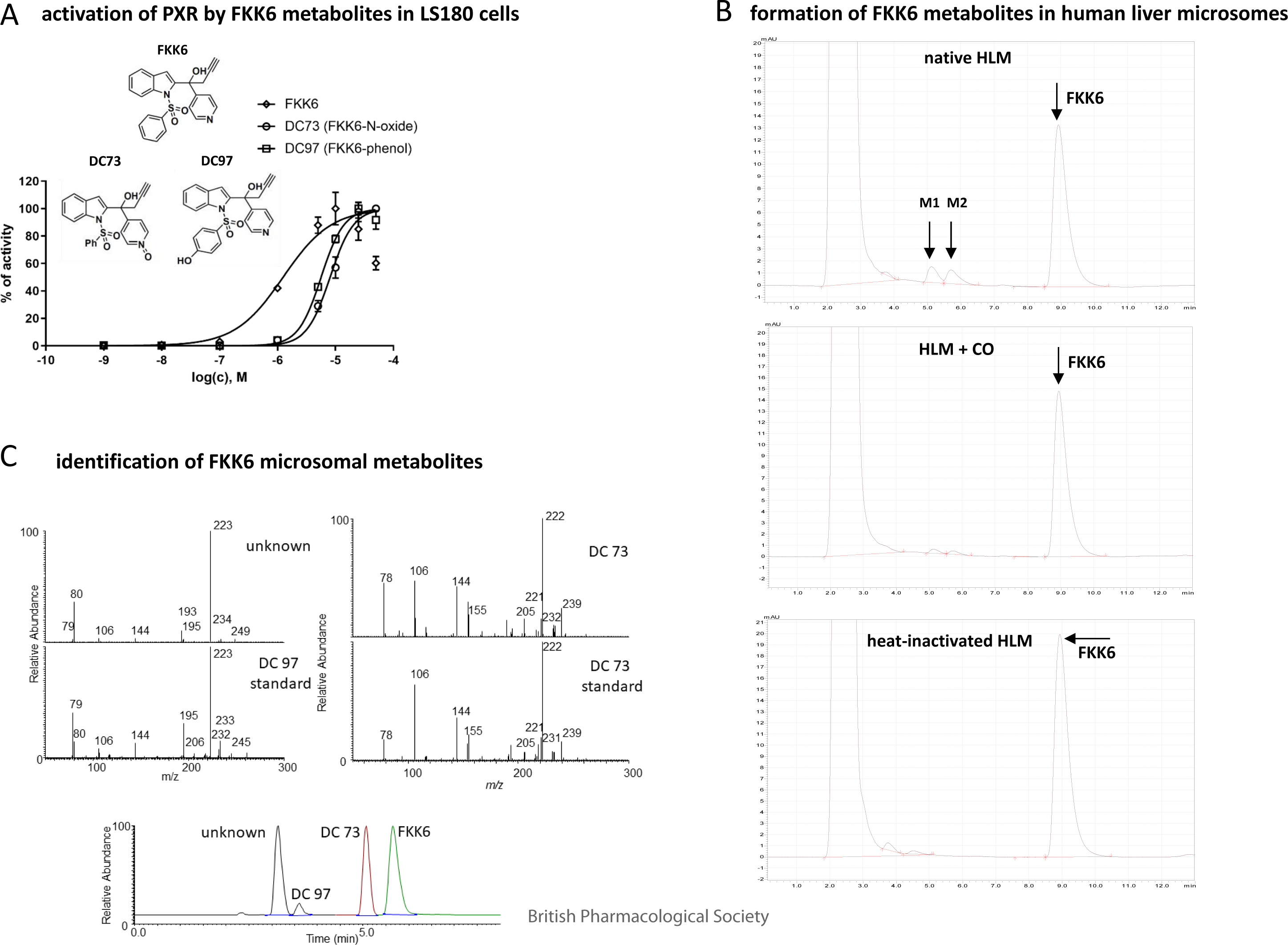
*In vitro* metabolism of FKK6. ***(A) Activation of PXR by predicted FKK6 metabolites.*** Intestinal LS180 cells transiently transfected with pSG5-hPXR expression plasmid along with a luciferase reporter plasmid p3A4-luc were incubated for 24 h with DC73, DC97, FKK6, rifampicin (RIF; 10 µM) and vehicle (DMSO; 0.1% V/V). Cells were lysed and luciferase activity was measured. Incubations were carried out in quadruplicates (technical replicates). The data show mean ± SD from the experiments performed in three consecutive cell passages, and they are expressed as a percentage of PXR activity achieved by RIF. ***(B,C) Metabolism of FKK6 in human liver microsomes.*** Human liver microsomes (HLM) were incubated for 30 min with FKK6 (50 µM), and UV-HPLC analyses were then carried out; upper panel – native HLM, middle panel – CO-saturated HLM, lower panel – heat-inactivated HLM (B). LC/MS chromatogram and mass spectra of FKK6 microsomal metabolites (C).

Finally, we identified the structures of FKK6 metabolites using LC/MS. Three mass spectrometric signals of putative FKK6 metabolites were acquired (Figure 4C) in samples from microsomal incubations of FKK6. The most polar metabolite (designated as unknown) shows lower retention than the authentic standard of DC97 but a similar fragmentation pattern and UV profile. The same spectral properties but different chromatographic behavior suggest that “unknown” might be an isomer of *in silico* predicted metabolite DC97 (C-19 phenol). The latter was found in trace amounts that did not allow for proper spectral identification. The third signal was assigned to DC73 (N6-oxide) based on co-elution with authentic standards and the same spectral properties. Quantification of DC73 was performed. In microsomal incubation mixtures with 10 µM and 50 µM FKK6, the concentrations of DC73 reached 0.98 ± 0.15 µM (n=3) and 0.66 ± 0.10 µM (n=2), respectively. The lack of correlation between the concentration of the parental compound and its metabolite indicates the mixed effect of FKK6 at CYP enzymes, i.e., being both substrate and inhibitor.

## DISCUSSION

FKK6 is a promising PXR-modulating candidate for locoregional rodent and human intestinal inflammation therapy. We have demonstrated previously that FKK6 is a modest PXR agonist ligand [12], and have argued for developing PXR modest agonists as potential new therapeutics for inflammatory bowel disease [8]. However, before proceeding with a preclinical development plan, we sought to clarify the basic in vitro pharmacologic properties of FKK6. As suspected, FKK6 exerts a weak conformational change and a distinct binding mode in contrast to strong agonists such as SR12813. The importance of this finding is that sufficiently high concentrations of FKK6 would be required to gain some level of PXR activation in inflamed tissues, and the same would be required if FKK6 were absorbed and retained essentially as a parent molecule. However, the in vitro metabolism screen for FKK6 suggests rapid metabolism through the CYP450 system and that the metabolites (at least two predicted forms) are far less active on PXR. However, it is to be noted that a major unknown metabolite has yet to be characterized.

Nevertheless, the rapid clearance of FKK6 may be a limiting factor for bioavailability, as it may undergo first-pass metabolism in the liver after oral dosing, which can limit the amount of drug reaching the circulation, even if the compound is well absorbed. Assuming idiosyncratic side-effects of the formed metabolites are absent, this attribute favors the use of FKK6 in locoregional delivery to the colon. The liver-based effects on PXR would likely be minimal.

FKK6, as synthesized, exists in a racemic mixture, and efforts are underway to purify its enantiomers [12]. It is reasonable to speculate that one of the enantiomers demonstrates significant binding, while the other does not. Moreover, the presence of various derivatives of FKK6 metabolites should not come as a surprise, as illustrated by the unidentified metabolite in Figure 4C. Other low-abundance metabolites would also likely exist. The plans are to purify and quantify existing metabolites from microsomal incubations and to validate their effects (or lack thereof) on PXR activation. In this context, in mice dosed with FKK6 via oral gavage, there is no significant PXR target gene induction (mdr1, cyp3a11) in the liver of mice (wild-type mice, mice expressing the human PXR transgene, and mice with endogenous mouse PXR knockout)[12]. While requiring further validation, these data may suggest that even mouse metabolites of FKK6 are not circulating in high concentrations to affect PXR activation in the liver. However, these results could be a function of the administered dose, and further studies are needed to validate these findings.

Indoles (and a variety of analogs) are known to bind and alter the GPCR signaling pathway [39, 40]. Similarly, indole compounds can also activate nuclear receptors other than the orphan receptors we tested in our previous manuscript [12, 41, 42]. Our extended panel of orphan and non-orphan GPCRs and steroid hormone (e.g., ER) and non-steroid hormone nuclear receptors (e.g., PPAR) demonstrated that FKK6 has minimal agonist effects. In the antagonist assay for the synthetic ligand-activated GPCRs and nuclear receptors, FKK6 induced > 50% antagonism of cannabinoid receptor 1 (CNR1), GPR103, GPR92, follicle-stimulating hormone receptor (FSHR) only, and the progesterone receptor (PR), respectively. The physiological relevance of this finding remains unclear and requires additional in vivo testing in relevant receptor model systems.

Overall, FKK6 is a promising candidate for the initiation of preclinical discovery approaches toward the amelioration of inflammatory bowel disease. It is cautioned that a significant amount of detailed work is required regarding pharmacology and toxicology for the compound to be tested in genetic and other models of IBD. These studies are underway, and appropriate formulations will also be conducted before validating the effects of FKK6 in preclinical models of IBD.

### Study limitations

Except *in vitro* metabolism studies, the studies presented were conducted *in vitro* at single FKK6 concentrations (10 μM) and more detailed testing using a range of concentrations of FKK6 will provide a clearer picture of the off-target effects of FKK6. However, in defense of this single concentration, all our *in vitro* and cell line reporter studies have been performed using 10 μM for PXR effects [12]. Since we do not have pharmacokinetic data, these concentrations must be validated *in vivo*. Detailed studies on metabolites in rodent and human systems will need to be clarified, especially concerning PXR activation. Additional off-target investigations are needed to decipher how FKK6 might work in cells. Despite these limitations, the current data suggests a safety signal worthy of further preclinical pursuit of a novel microbial metabolite mimic for IBD.

## ACKNOWLEDGEMENTS

Financial support from Czech Science Foundation grant #22-00355S (to Z.D.) is greatly acknowledged. We thank dr. Ján Vančo for determination of the partition coefficient.

## AUTHORŚ CONTRIBUTIONS

CrediT STATEMENT: Conceptualization**^1^**, Data Curation**^2^**, Formal Analysis**^3^**, Funding Acquisition**^4^**, Investigation**^5^**, Methodology**^6^**, Project Administration**^7^**, Resources**^8^**, Supervision**^9^**, Validation**^10^**, Visualization**^11^**, Writing – Original Draft Preparation**^12^**, Writing – Review & Editing**^13^**

Zdeněk Dvořák^1,3,4,5,7,8,9,11,12,13^, Barbora Vyhlídalová^3,7,11^, Petra Pečinková^2,3,5^, Hao Li^2,3,5^, Pavel Anzenbacher^2,3,5,6,9,10,11^, Alena Špičáková^2,3,5,6^, Eva Anzenbacherová^2,3,5,6^, Vimanda _Chow2,3,5,6,10,11, Jiabao Liu2,3,5,6,10,11, Henry Krause2,3,5,6,10,11, Derek Wilson2,3,5,6,10,11, Tibor Berés2,3,5,6,10,11, Petr Tarkowski2,3,5,6,10,11, Dajun Chen2,3,5,6,10,11, Sridhar Mani1,3,5,7,8,9,10,11,12,13_

## DATA AVAILABILITY STATEMENT

The data that support the findings of this study are openly available in ZENODO at http://doi.org/10.5281/zenodo.10008305.

## ABBREVIATIONS

AR: Androgen Receptor
BCRP: Breast Cancer Resistance Protein
BSEP: Bile Salt Export Pump
EFC: Enzyme Fragment Complementation
ERα: Estrogen Receptor α
FXR: Farnesoid X Receptor
GPCR: G-Protein Coupled Receptor
GR: Glucocorticoid Receptor
hCa_V_1.2h: Human Calcium Ion Channel
hK_V_11.1. hERG: Human Potassium Ion Channel
Na_V_1.5: Human Sodium Ion Channel
HLM: Human Liver, Microsomes
LXRα/β: Liver X receptor α/β
MATE: Multidrug And Toxin Extrusion protein
MR: Mineralocorticoid Receptor
MRP2: Multidrug Resistance-associated Protein
OAT: Organic Anion Transporter
OATP: Organic Anion Transporting Polypeptide
OCT: Organic Cation Transporter
PPARα/δ/γ: Peroxisome Proliferator-Activated Receptor α/δ/γ
PR A/B: Progesterone Receptor A/B
P-gp: Multidrug Resistance Protein 1
RARα/β: Retinoic Acid Receptor α/β
RXRα/γ: Retinoid X Receptor α/γ
SRCP: Steroid Receptor Co-activator Peptide
THRα/β: Thyroid Hormone Receptor α/β

## FIGURE LEGEND

**Figure S1.**
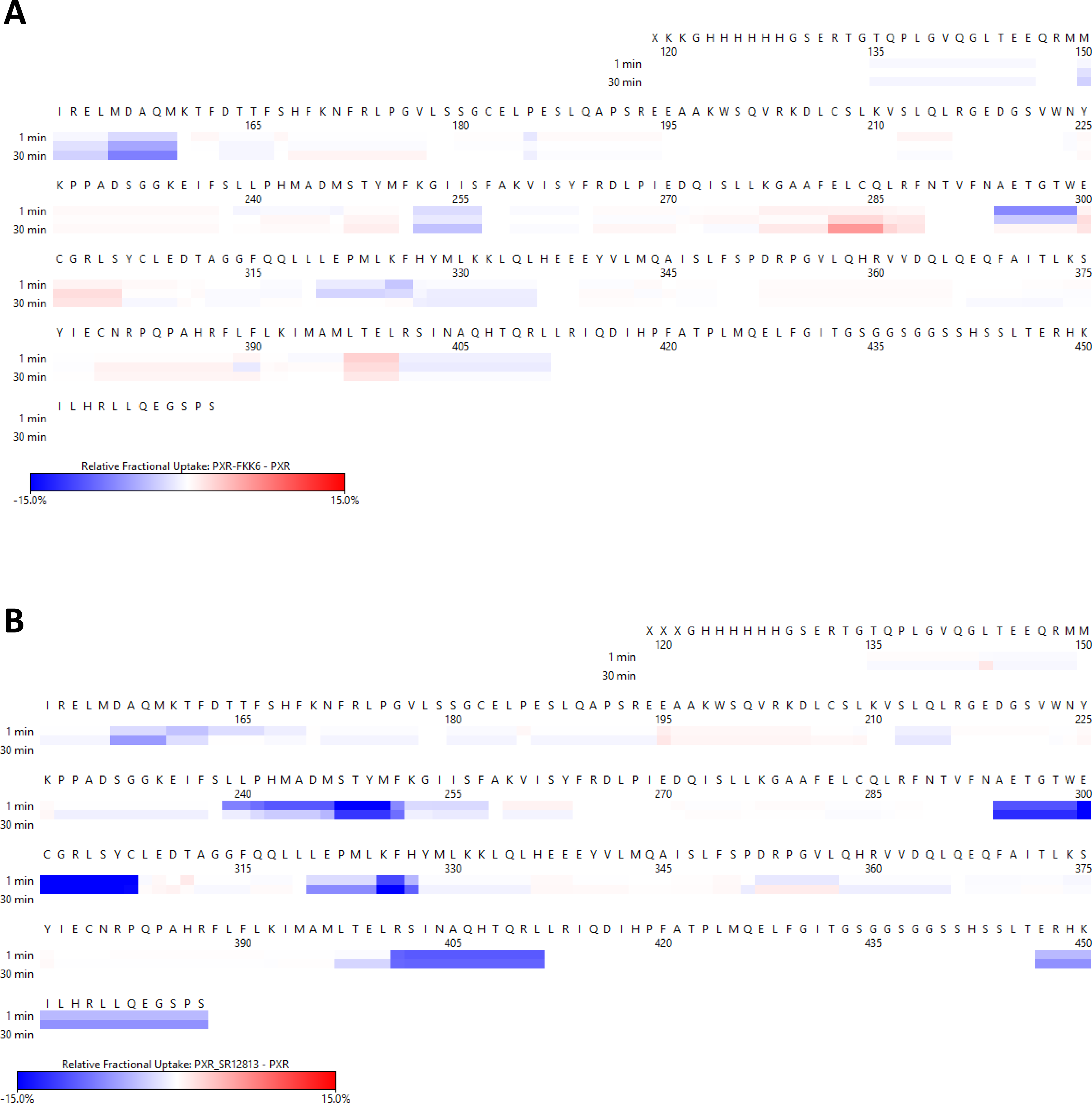
Heat map for the differences in deuterium uptake. Time intervals at 1, 10 and 30 min between ***(A)*** Apo PXR and PXR-bound SR12813 and ***(B)*** Apo PXR and PXR-bound FKK6.

**Figure S2.**
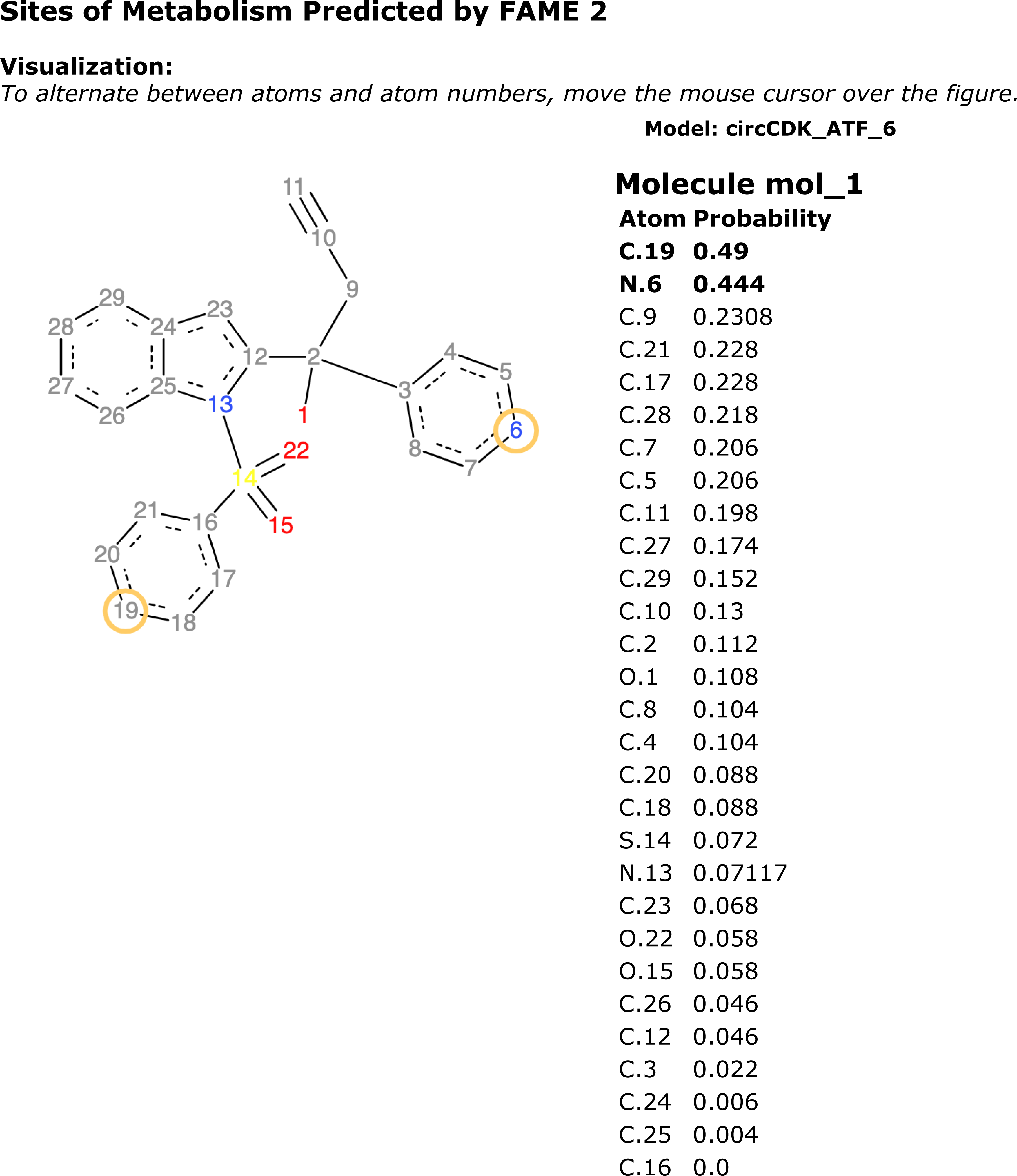
Predicted sites of metabolism of FKK6 by FAME2.

## REFERENCES

1. Vázquez-Gómez, G., et al., Aryl hydrocarbon receptor (AhR) and pregnane X receptor (PXR) play both distinct and common roles in the regulation of colon homeostasis and intestinal carcinogenesis. Biochem Pharmacol, 2023. 216: p. 115797.

2. Sun, L., et al., Role of nuclear receptor PXR in immune cells and inflammatory diseases. Front Immunol, 2022. 13: p. 969399.

3. Bautista-Olivier, C.D. and G. Elizondo, PXR as the tipping point between innate immune response, microbial infections, and drug metabolism. Biochem Pharmacol, 2022. 202: p. 115147.

4. Klepsch, V., et al., Nuclear Receptors Regulate Intestinal Inflammation in the Context of IBD. Front Immunol, 2019. 10: p. 1070.

5. Mackowiak, B., et al., The Roles of Xenobiotic Receptors: Beyond Chemical Disposition. Drug Metab Dispos, 2018. 46(9): p. 1361–1371.

6. Cheng, J., Y.M. Shah, and F.J. Gonzalez, Pregnane X receptor as a target for treatment of inflammatory bowel disorders. Trends Pharmacol Sci, 2012. 33(6): p. 323–30.

7. Kliewer, S.A., B. Goodwin, and T.M. Willson, The nuclear pregnane X receptor: a key regulator of xenobiotic metabolism. Endocr Rev, 2002. 23(5): p. 687–702.

8. Dvořák, Z., H. Li, and S. Mani, Microbial Metabolites as Ligands to Xenobiotic Receptors: Chemical Mimicry as Potential Drugs of the Future. Drug Metab Dispos, 2023. 51(2): p. 219–227.

9. Dvořák, Z., H. Sokol, and S. Mani, Drug Mimicry: Promiscuous Receptors PXR and AhR, and Microbial Metabolite Interactions in the Intestine. Trends Pharmacol Sci, 2020. 41(12): p. 900–908.

10. Sartor, R.B., Review article: the potential mechanisms of action of rifaximin in the management of inflammatory bowel diseases. Aliment Pharmacol Ther, 2016. 43 **Suppl 1**: p. 27–36.

11. Kumar, J. and A.M.J. Newton, Rifaximin - Chitosan Nanoparticles for Inflammatory Bowel Disease (IBD). Recent Pat Inflamm Allergy Drug Discov, 2017. 11(1): p. 41–52.

12. Dvorak, Z., et al., Targeting the pregnane X receptor using microbial metabolite mimicry. EMBO Mol Med, 2020. 12(4): p. e11621.

13. Dvorak, Z., H. Sokol, and S. Mani, Drug Mimicry: Promiscuous Receptors PXR and AhR, and Microbial Metabolite Interactions in the Intestine. Trends Pharmacol Sci, 2020. 41(12): p. 900–908.

14. Vyhlidalova, B., et al., Differential activation of human pregnane X receptor PXR by isomeric mono-methylated indoles in intestinal and hepatic in vitro models. Toxicol Lett, 2020. 324: p. 104–110.

15. Vyhlidalova, B., et al., Mono-methylindoles induce CYP1A genes and inhibit CYP1A1 enzyme activity in human hepatocytes and HepaRG cells. Toxicol Lett, 2019. 313: p. 66–76.

16. Stepankova, M., et al., Methylindoles and Methoxyindoles are Agonists and Antagonists of Human Aryl Hydrocarbon Receptor. Molecular Pharmacology, 2018. 93(6): p. 631–644.

17. Illés, P., et al., Indole microbial intestinal metabolites expand the repertoire of ligands and agonists of the human pregnane X receptor. Toxicology Letters, 2020. 334: p. 87–93.

18. Vyhlidalova, B., et al., Gut Microbial Catabolites of Tryptophan Are Ligands and Agonists of the Aryl Hydrocarbon Receptor: A Detailed Characterization. Int J Mol Sci, 2020. 21(7).

19. Sládeková, L., et al., Switching on/off aryl hydrocarbon receptor and pregnane X receptor activities by chemically modified tryptamines. Toxicol Lett, 2023. 387: p. 63–75.

20. Li, H., et al., Deciphering structural bases of intestinal and hepatic selectivity in targeting pregnane X receptor with indole-based microbial mimics. Bioorg Chem, 2021. 109: p. 104661.

21. Liu, J., et al., The omega-3 hydroxy fatty acid 7(S)-HDHA is a high-affinity PPARalpha ligand that regulates brain neuronal morphology. Sci Signal, 2022. 15(741): p. eabo1857.

22. Dierks, E.A., et al., A method for the simultaneous evaluation of the activities of seven major human drug-metabolizing cytochrome P450s using an in vitro cocktail of probe substrates and fast gradient liquid chromatography tandem mass spectrometry. Drug Metab Dispos, 2001. 29(1): p. 23–9.

23. Zhang, L., et al., Cloning and functional expression of a human liver organic cation transporter. Mol Pharmacol, 1997. 51(6): p. 913–21.

24. Kido, Y., P. Matsson, and K.M. Giacomini, Profiling of a prescription drug library for potential renal drug-drug interactions mediated by the organic cation transporter 2. J Med Chem, 2011. 54(13): p. 4548–58.

25. Cihlar, T. and E.S. Ho, Fluorescence-based assay for the interaction of small molecules with the human renal organic anion transporter 1. Anal Biochem, 2000. 283(1): p. 49–55.

26. Gui, C., et al., Development of a cell-based high-throughput assay to screen for inhibitors of organic anion transporting polypeptides 1B1 and 1B3. Curr Chem Genomics, 2010. 4: p. 1–8.

27. Yasujima, T., et al., Evaluation of 4’,6-diamidino-2-phenylindole as a fluorescent probe substrate for rapid assays of the functionality of human multidrug and toxin extrusion proteins. Drug Metab Dispos, 2010. 38(4): p. 715–21.

28. Paturi, D.K., et al., Identification and functional characterization of breast cancer resistance protein in human bronchial epithelial cells (Calu-3). Int J Pharm, 2010. 384(1-2): p. 32–8.

29. Matsson, P., et al., Identification of novel specific and general inhibitors of the three major human ATP-binding cassette transporters P-gp, BCRP and MRP2 among registered drugs. Pharm Res, 2009. 26(8): p. 1816–31.

30. Polli, J.W., et al., Rational use of in vitro P-glycoprotein assays in drug discovery. J Pharmacol Exp Ther, 2001. 299(2): p. 620–8.

31. Dawson, S., et al., In vitro inhibition of the bile salt export pump correlates with risk of cholestatic drug-induced liver injury in humans. Drug Metab Dispos, 2012. 40(1): p. 130–8.

32. Lipinski, C.A., et al., Experimental and computational approaches to estimate solubility and permeability in drug discovery and development settings. Adv Drug Deliv Rev, 2001. 46(1-3): p. 3–26.

33. Banker, M.J., T.H. Clark, and J.A. Williams, Development and validation of a 96-well equilibrium dialysis apparatus for measuring plasma protein binding. J Pharm Sci, 2003. 92(5): p. 967–74.

34. Obach, R.S., et al., The prediction of human pharmacokinetic parameters from preclinical and in vitro metabolism data. J Pharmacol Exp Ther, 1997. 283(1): p. 46–58.

35. Suzuki, A., et al., Identification of human cytochrome P-450 isoforms involved in metabolism of R(+)- and S(-)-gallopamil: utility of in vitro disappearance rate. Drug Metab Dispos, 1999. 27(11): p. 1254–9.

36. Hidalgo, I.J., T.J. Raub, and R.T. Borchardt, Characterization of the human colon carcinoma cell line (Caco-2) as a model system for intestinal epithelial permeability. Gastroenterology, 1989. 96(3): p. 736–49.

37. Sicho, M., et al., FAME 2: Simple and Effective Machine Learning Model of Cytochrome P450 Regioselectivity. J Chem Inf Model, 2017. 57(8): p. 1832–1846.

38. Dong, X., et al., Copper-catalyzed trifluoromethylation and cyclization of aromatic-sulfonyl-group-tethered alkenes for the construction of 1,2-benzothiazinane dioxide type compounds. Chemistry, 2013. 19(50): p. 16910–5.

39. Bhattarai, Y., et al., Gut Microbiota-Produced Tryptamine Activates an Epithelial G-Protein-Coupled Receptor to Increase Colonic Secretion. Cell Host Microbe, 2018. 23(6): p. 775–785.e5.

40. Obi, P. and S. Natesan, Membrane Lipids Are an Integral Part of Transmembrane Allosteric Sites in GPCRs: A Case Study of Cannabinoid CB1 Receptor Bound to a Negative Allosteric Modulator, ORG27569, and Analogs. J Med Chem, 2022. 65(18): p. 12240–12255.

41. Darwish, K.M., et al., Synthesis, biological evaluation, and molecular docking investigation of benzhydrol- and indole-based dual PPAR-γ/FFAR1 agonists. Bioorg Med Chem Lett, 2018. 28(9): p. 1595–1602.

42. Zhu, Y., et al., Research progress of indole compounds with potential antidiabetic activity. Eur J Med Chem, 2021. 223: p. 113665.

